# The impact of environmental exposures on the epigenomic and transcriptomic landscape of transposable elements

**DOI:** 10.1101/2025.07.28.667212

**Authors:** Benpeng Miao, Yu Zhang, Wanqing Shao, Shuhua Fu, Marisa S. Bartolomei, Cheryl Walker, Cristian Coarfa, TaRGET II Consortium, Bo A. Zhang, Ting Wang

**Affiliations:** Department of Genetics, Washington University School of Medicine, St. Louis, MO 63108, USA; Department of Developmental Biology, Washington University School of Medicine, St. Louis, MO 63108, USA; Center of Regenerative Medicine, Washington University School of Medicine, St. Louis, MO 63108, USA; Department of Cell and Developmental Biology, Perelman School of Medicine, University of Pennsylvania, Philadelphia, PA 19104, USA; Dan L. Duncan Comprehensive Cancer Center, Baylor College of Medicine, Houston, TX 77030, USA; Center for Precision Environmental Health, Baylor College of Medicine, Houston, TX 77030, USA

**Keywords:** Transposable elements, environmental exposures, regulatory regions, transcriptome diversity

## Abstract

Transposable elements (TEs) are mobile DNA sequences that constitute a significant portion of mammalian genomes. While typically silenced by epigenetic mechanisms, mounting evidence indicates TEs can regulate gene expression and chromatin architecture. However, their regulatory roles under various environmental exposures remain largely unexplored. In this study, we investigate the regulatory functions of TEs in mouse liver tissue following early-life exposure to environmental toxicants, including arsenic (As), lead (Pb), bisphenol A (BPA), tributyltin (TBT), di-2-ethylhexyl phthalate (DEHP), tetrachlorodibenzo-p-dioxin (TCDD), and particulate matter less than 2.5 micrometers (PM2.5). These toxicants are linked to various health issues, including neurodevelopmental deficits, metabolic and immune dysfunction, and increased cancer risks. Integrative analysis of 351 multi-omics datasets from liver tissues of 5-month-old mice indicated that early-life environmental exposures significantly altered chromatin accessibility and expression of TEs in later life stage, revealing distinct exposure-specific signatures and sex-dependent responses. 6,699 TEs were identified with altered chromatin accessibility, mostly in non-coding regions, suggesting potential impact on gene regulation. Within these TEs, LINE elements were enriched in genes involved in metabolic pathways, while LTR elements, particularly the ORR1E subfamily, were predominantly associated with immune-related genes. Additionally, we identified 140 TE-gene chimeric transcripts with TE-derived novel transcription start sites, highlighting TE-contributed transcriptional plasticity. Our findings depict a comprehensive landscape of TE regulation under early-life toxicant exposures, offering insights into TEs biology and their impact on health and disease.

## Introduction

Transposable elements (TEs) are repetitive DNA sequences with the capability to move within genomic DNA. Since discovered by Barbara McClintock in maize, TEs have been found in nearly all eukaryotic genomes, comprising up to 45% of the human genome and ∼40% of the mouse genome^1,2^. TEs are commonly classified into two primary categories: DNA transposons, which mobilize themselves in the genome through a DNA-mediated “cut-and-paste” mechanism, and retrotransposons, including LINE, SINE, and LTR, which replicate through an RNA intermediate “copy and paste” mechanism, resulting in potentially hundreds of thousands of copies in the genome^1,2^. Although historically dismissed as “junk DNA” or selfish genetic elements, mounting evidence supports the substantial roles of TEs in genome regulation, evolution, and adaptation^3–6^. TEs can contribute to genomic rearrangements and lead to significant genome recombination^7,8^. Through replication in the host genome, TEs act as sources of regulatory sequences, such as promoters, enhancers, repressors, and insulators, which can regulate gene expression and reshape the epigenetic landscap^9–13^. Many TEs also harbor transcription factor binding sites, and their insertion into new genomic contexts can create or modify cis-regulatory elements^6,14,15^, facilitating the formation of new transcriptional variants through distinct mechanisms, such as the creation of novel transcription start or termination sites, exon shuffling, and alternative splicing^16–19^. These mechanisms enable TEs to generate genetic diversity and potentially create new functional elements within the host genome, facilitating adaptation to changing environmental conditions.

Under normal conditions, transposable elements (TEs) are typically repressed by epigenetic mechanisms, including DNA methylation and histone modifications, to preserve genome stability and prevent aberrant activation^20–23^. However, environmental stimuli, particularly exposure to heavy metals, endocrine-disrupting chemicals, and particulate matter, can trigger widespread epigenetic changes that disrupt TE repression and activate cellular stress responses^24,25^. Exposure to environmental toxicants is widely recognized as a serious threat to human health. Arsenic and lead are toxic heavy metals linked to neurological and cardiovascular disorders^26,27^. BPA and DEHP, commonly used in plastic production, are endocrine disruptors that interfere with hormonal signaling and increase metabolic disorders and cancer risks^28,29^. TBT, a toxic organotin compound used in antifouling paints and wood preservatives, is among the most damaging pollutants in aquatic environments and poses significant risks to human health^30^. TCDD, a persistent organic pollutant, is associated with reproductive, developmental, and hormonal disturbances^31^. PM2.5, a major component of air pollution, contributes to respiratory and cardiovascular diseases and is associated with a wide range of health complications^32,33^.

Exposure to these pollutants has long-lasting effects and significantly elevates disease risk. Studies have shown that such exposures can epigenetically deregulate TEs^24^. For instance, Exposure to carcinogen benzo(a)pyrene (BaP) increases LINE-1 retrotransposition in HeLa cells through genotoxic stress and epigenetic remodeling, including loss of DNA methylation and the increase of active histone marks^34,35^. Similarly, low-level arsenic exposure results in L1 retrotransposition in HepG2 cells^36^. Meanwhile, *in vitro* and epidemiological studies suggest that the DNA methylation level of TEs is associated with arsenic and lead exposure^37,38^. Dolinoy *et al.* demonstrated that maternal exposure to BPA in *Agouti* viable yellow (Avy) mouse model can lead to hypomethylation of an IAP retrotransposon upstream of the Agouti gene, resulting in a shift in offspring coat color toward yellow^39^. These findings emphasized that environmental stressors can epigenetically alter transposable element regulation, particularly through changes in DNA methylation, which may contribute to adverse developmental and physiological outcomes.

In this study, we collected liver transcriptome and epigenome data from 5-month-old mice, which were exposed to toxic pollutants during early development, to systematically investigate how TEs respond to early-life exposure of toxic environmental pollutants, including As, Pb, BPA, TBT DEHP, TCDD, and two types of PM2.5 generated by University of Chicago (PM2.5-CHI) and Johns Hopkins University (PM2.5-JHU) with distinct components. Our analysis revealed exposure-specific and sex-specific patterns of TEs in mouse liver tissue, including alterations in chromatin accessibility and expression in response to environmental toxicants. In total, 6,699 TEs, most of them located in non-coding cis-regulatory regions, significantly altered chromatin accessibility in response to at least one toxicant exposure. Genes around distinct classes of TEs were associated with different biological processes: genes around LINEs were associated with metabolic processes, while genes around TEs from the LTR class were enriched in immune-related functions, especially those from the ORR1E subfamily. Meanwhile, 540 TE-derived known transcripts and 140 TE-derived novel TE-gene chimeric transcripts were significantly dysregulated by at least one toxicant exposure, and most of them were upregulated, highlighting the reactivation of TE under stimulus. Our analysis charted the landscape of active regulation of TEs and highlighted their crucial roles in gene expression regulation under environmental stress.

## Results

### Exposures induced expression and accessibility changes of TE subfamilies

To explore the roles of transposable elements in mice responding to environmental exposures, we collected a total of 178 RNA-seq and 173 ATAC-seq datasets of liver samples from 5-month-old mice in the TaRGET II consortium^40^. The mice were exposed to As, Pb, BPA, TBT, DEHP, TCDD, or two distinct types of air pollution (PM2.5-CHI and PM2.5-JHU) two weeks prior to conception, through gestation and weaning. The exposures were withdrawn at postnatal 3 weeks, and the mice were maintained in exposure-free condition until 5 months (Fig. 1a and Supplementary file 1). The transcriptomes (RNA-seq) and epigenomes (ATAC-seq) of livers were profiled in both exposed and control groups, to characterize the molecular changes induced by early-life exposure.

**Figure 1.**
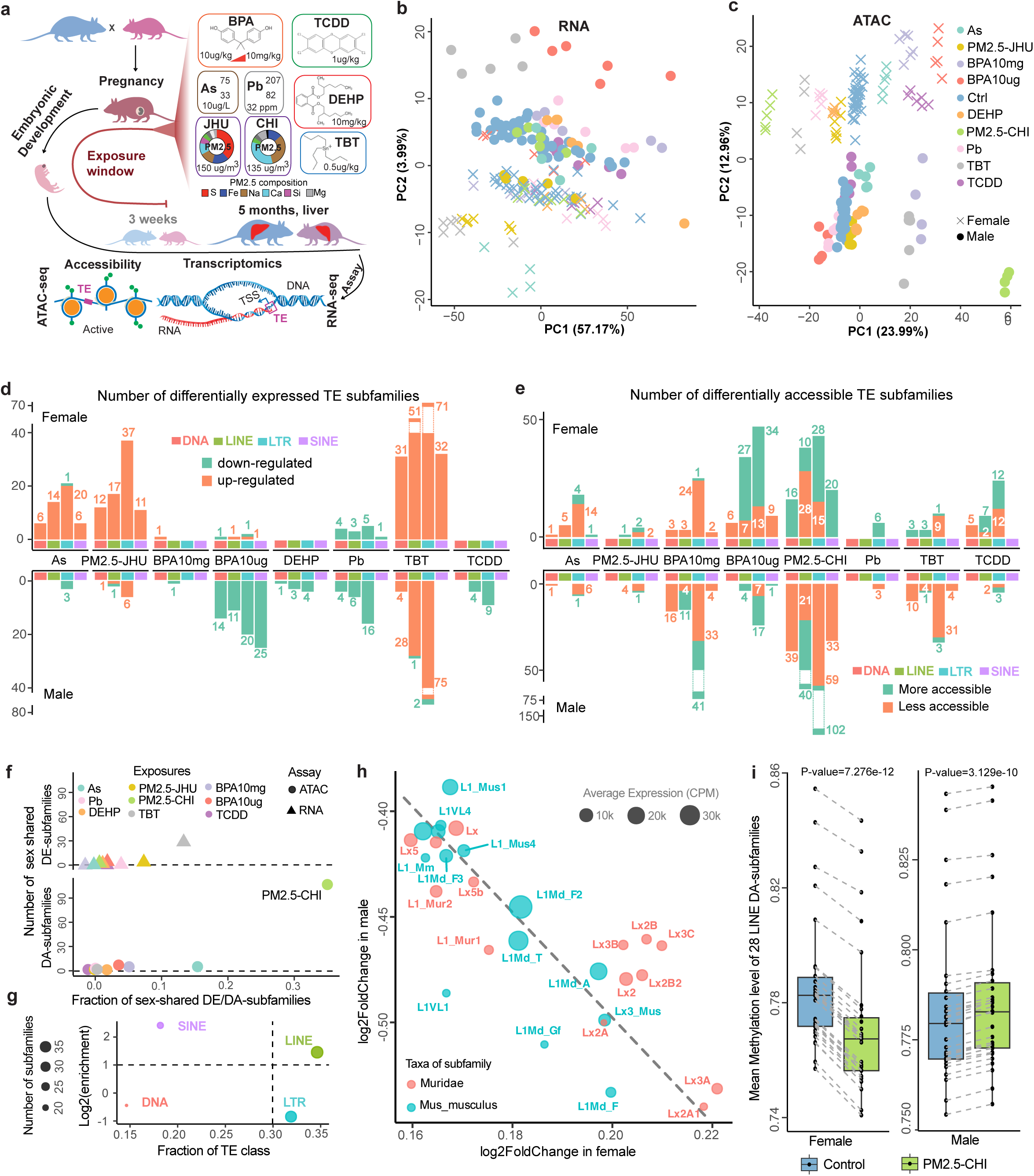
Exposures-induced expression and chromatin accessibility changes of transposable elements (TE) at the subfamily level. **a**) Study design: The transcriptome and epigenome of the 5-month-old mouse livers, which were exposed to different environmental toxicants at the early-life stage. As: arsenic; Pb: lead; BPA: bisphenol A; TBT: tributyltin; DEHP: the phthalate di-2-ethylhexyl phthalate; TCDD: the dioxin tetrachlorodibenzo-p-dioxin (TCDD); PM2.5: air pollution in the form of particulate matter <2.5μm. CHI: University of Chicago; JHU: Johns Hopkins University. S: sulfur; Fe: iron; Na: sodium; Ca: calcium; Si: silicon; Mg: magnesium. The PCA plot for RNA-seq **b**) and ATAC-seq **c**) displayed the distribution of samples under distinct exposures (color-coded). Dot: male sample; cross: female sample. **d**) Number of up-(orange) and down-regulated (green) differentially expressed TE subfamilies (DE-subfamilies) in females and males under different exposures. Largest number of DE-subfamilies was identified in TBT-exposed livers. **e**) Number of more accessible (orange) and less accessible (green) differentially accessible TE subfamilies (DA-subfamilies) under different exposures. **f**) Number of DE- and DA-subfamilies that were shared between female and male. Very few sex-shared DE- and DA-subfamilies were found in most of the exposures. **g**) Enrichment of TE classes of DA-subfamilies shared between female and male under the PM2.5-CHI. Circle size: number of DA-subfamilies in different TE classes. **h**) Accessibility changes of young LINE DA-subfamilies shared between PM2.5-CHI female and male. These subfamilies became more accessible in females but less accessible in males following PM2.5-CHI exposure. Red and green: taxa of TE subfamilies. Dot size: normalized expression in control and PM2.5-CHI samples. **i**) Significant hypo-methylation of 28 LINE DA-subfamilies in PM2.5-CHI, when compared to control samples (paired Wilcoxon signed rank exact test).

We first explored the transcriptomic and epigenomic changes of TEs under exposure at the subfamily level. There were no obvious global changes of expression or chromatin accessibility in TE subfamilies under exposure conditions when compared with controls (S-fig. 1a). However, the principal component analysis (PCA) strongly indicated that the distinct long-term system-wide effects induced by early-life exposures were highly sex-specific and toxicant-dependent (Fig. 1b and c). At the transcriptional level, PM2.5-JHU and TBT exposure induced significant changes in both sexes, whereas other exposures had only minor effects, except for BPA10μg in male liver and As in female liver, showing notable transcriptional responses (Fig.1b). A clearer pattern can be observed at the chromatin accessibility level. Female livers showed diverse chromatin accessibility changes in response to various exposures, whereas male livers exhibited significant disruption only after exposure to PM2.5-CHI, TBT, and BPA10mg (Fig. 1c). We then identified significantly differentially expressed TE subfamilies (DE-subfamilies) (Fig. 1d and Supplementary file 2) and differentially accessible TE subfamilies (DA-subfamilies) (Fig. 1e and Supplementary file 3) in different exposure conditions, and the results aligned well with PC analysis. For example, TBT exposure significantly up-regulated the expression of DE-subfamilies in both sexes. We identified more up-regulated DE-subfamilies in females exposed to As and PM2.5-JHU, whereas BPA10ug, DEHP, Pb, and TCDD induced more male-specific down-regulated DE-subfamilies. Interestingly, more than half of the Muridae-specific TE subfamilies, which are the youngest in mouse genome evolution, especially SINE and LTR elements, were found to be highly responsive to the early-life toxicant exposures, when compared to the genome background in both sexes (S-fig. 1b). A similar pattern also emerged in DA-subfamilies analysis (S-fig. 1c).

We further investigated the sex-specific responses of transposable elements to early-life exposures and observed distinct patterns between males and females. Most of exposures induced sex-specific transcriptional and epigenetic changes at TE subfamily level (Fig. 1f and S-fig. 1d). Notably, early-life exposure to PM2.5-CHI led to opposite chromatin accessibility changes in 109 shared TE subfamilies between female and male mice (correlation coefficient = –0.595, S-fig. 1d and Supplementary file 4). Of these, LINE subfamilies made up 35% of the total, and 31% were LTR subfamilies (Fig. 1g and Supplementary file 5). We identified 28 out of 38 differentially accessible LINE subfamilies as Muridae-specific young elements, all of which became more accessible in females but less accessible in males following PM2.5-CHI exposure (Fig. 1h and Supplementary file 6). This sex-biased chromatin accessibility was consistent with DNA methylation alteration patterns in matched samples: these 28 LINE subfamilies were generally hypomethylated in PM2.5-CHI females and hypermethylated in PM2.5-CHI males compared to controls (Fig. 1i).

### Epigenetic changes of transposable elements in response to environmental exposures

To investigate the potential regulatory role of transposable elements in response to environmental exposures, we identified the open chromatin regions (OCRs) derived from TEs (OCR-TEs) by analyzing ATAC-seq data across all samples. OCR-TEs were defined as transposable elements that harbored ATAC-seq peak summits (**Methods**). The proportion of OCR-TEs relative to total OCRs was similar between exposed and control groups, averaging around 20% (Fig. 2a). We then quantified the proportion of differentially accessible regions (DARs) derived from TEs (DAR-TEs) across TE classes and compared them to the OCR-TEs proportion in each exposure condition (S-fig. 2a). Notably, the fractions of DAR-TEs varied across all four major TE classes, with the greatest variability observed in SINEs, suggesting sex-specific patterns in a toxicant-dependent fashion. For example, ∼27% DAR-SINEs in male liver changed chromatin accessibility under Pb exposure, but BPA10mg only induced 12% DAR-SINEs alterations in male liver (S-fig. 2a).

**Figure 2.**
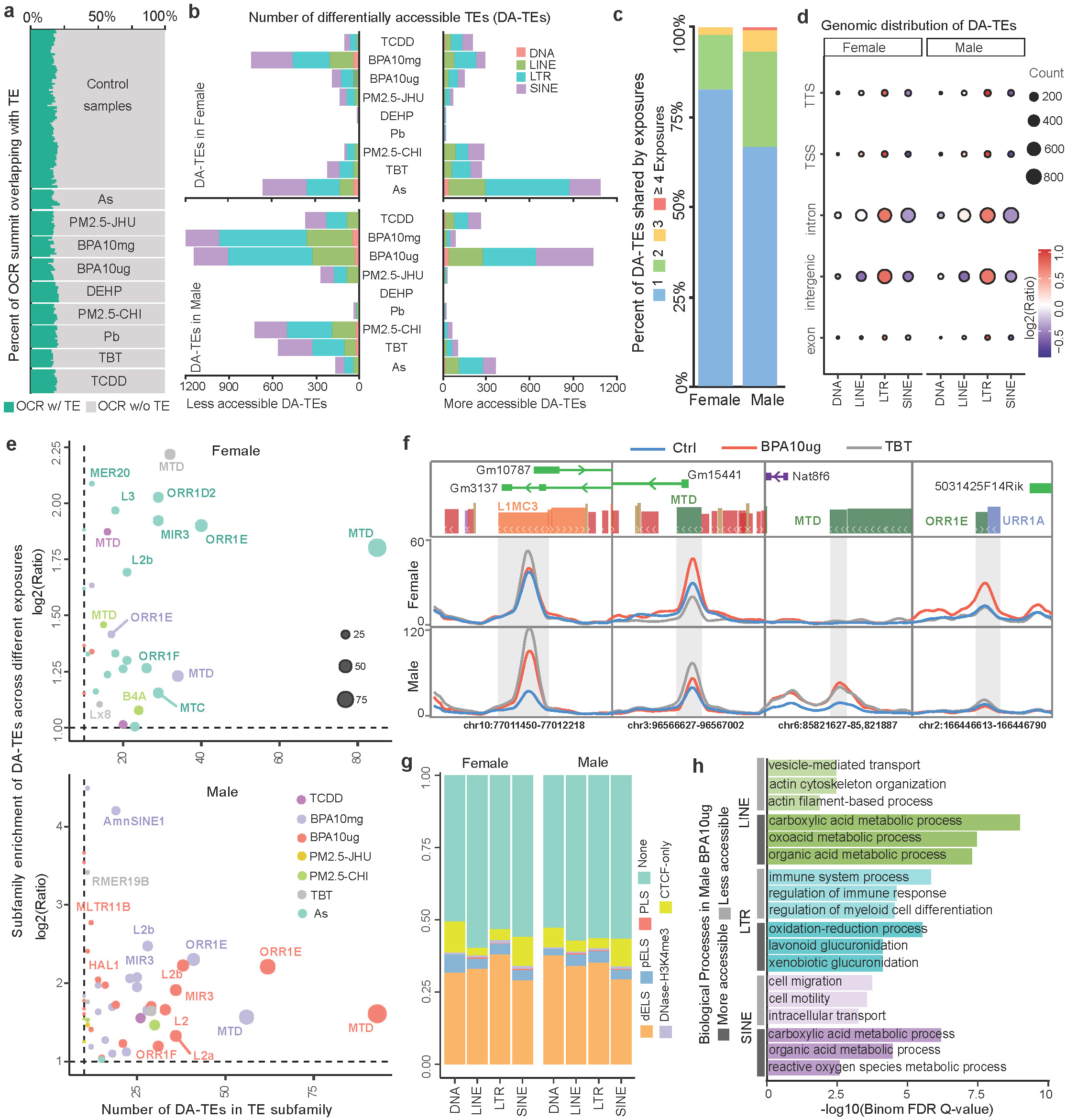
Chromatin accessibility changes of TE-derived regulatory elements under environmental exposures. **a**) ∼20% of open chromatin regions (OCRs) were derived from TEs across exposure and control samples. **b**) Number of differentially accessible TEs (DA-TEs) under different exposures. **c**) Percentages of DA-TEs identified in a single exposure or shared by multiple exposures separately for females and males. **d**) Genomic distribution enrichment of DA-TEs from different TE classes. The dot size was normalized to the count of DA-TEs. TSS: transcription start site; TTS: transcription terminate site. **e**) Subfamily enrichment of DA-TEs in female (top) and male (bottom) livers in response to environmental exposures. X-axis: the number of DA-TEs in each TE subfamily. Y-axis: log2 enrichment ratio to TE background in the mouse genome. Higher values indicate greater subfamily enrichment among DA-TEs. Dot size: number of DA-TEs in each subfamily. **f**) DA-TEs showing changed ATAC-seq signal in response to TBT and BPA10ug exposures compared to control. **g**) Fraction of DA-TEs with cis-regulatory elements annotation, including CTCF-only, DNase-H3K4me3, promoter (PLS), proximal and distal enhancer (pELS and dELS). None: DA-TEs without interaction between cis-regulatory elements. **h**) Enriched biological processes in DA-TEs of males in BPA10ug exposure.

We further characterized thousands of differentially accessible TEs (DA-TEs) that were identified in mouse livers under different exposures (**Methods**, Fig. 2b and Supplementary file 7). LINE and LTR classes of DA-TEs made up large proportions in most exposures, while limited DA-TEs were from DNA class (Fig. 2b). When comparing DA-TEs across different exposures, we found that 81% in female livers and 65% in male livers were unique to a single exposure, indicating that toxicant-induced DA-TEs had strong exposure specificity (Fig. 2c). The genomic distribution analysis showed that ∼80% of DA-TEs were located in the intergenic and intronic regions (S-fig. 2b), and LTR elements were over-represented in all genomic features when comparing to other TE classes, suggesting high activity potential of LTR elements in the mouse genome (Fig. 2d). We calculated the subfamily-enrichment of the DA-TEs in each exposure, and found that MTD enriched in multiple exposures in both female and male and ORR1E subfamilies were particularly enriched in As-exposed female livers and two BPA-exposed male livers (Fig. 2d and Supplementary file 8). These examples further demonstrated the dynamic responses of liver under different exposures (Fig. 2f and S-fig. 2d).

To explore the potential regulatory functions of DA-TEs, we examined cis-regulatory elements (cCREs) annotation from mouse ENCODE, and found that nearly half of DA-TEs were annotated as dELS and pELS enhancer elements (Fig. 2g and S-fig. 2c). The remaining half did not overlap with known cCREs and could be potential novel regulatory regions in responding to toxicant exposures. Interestingly, GO enrichment analysis for genes associated with DA-TEs revealed TE class-specific enrichment patterns (Supplementary file 9). LTRs were mainly enriched in immune processes, and LINE and SINEs were highly associated with metabolic, transport and cell organization (S-fig. 2e). For example, in males exposed to BPA10ug, genes associated with more accessible DA-LINEs and DA-SINEs were highly enriched in metabolic process, DA-LTRs around immune response-related genes tended to be closed under the exposure, and genes around less-opened DA-LINEs and DA-SINEs were more enriched in cell migration and actin organization (Fig. 2h).

We also observed a sex-specific pattern in DA-TE dysregulation (S-fig. 3a-c). In male livers, LTRs linked to immune-related processes showed altered chromatin accessibility exclusively in response to BPA exposure. In contrast, female livers exhibited changes in the accessibility of immune-associated LTRs specifically in response to As exposure (Fig. 3a). Further examination of TE subfamily enrichment of DA-LTRs associated with immune processes indicated ORR1E subfamily, most abundant DA-LTRs, was enriched (> 2-fold) for the immune process in responding to exposure of BPA and As (Fig. 3b and Supplementary file 10), but not enriched in metabolic process (S-fig. 3e). We identified 20 immune-associated ORR1E copies that changed chromatin accessibility (DA-ORR1Es), 8 of which responded to both BPA and As exposure (Fig. 3b), and they were associated with genes including *Naip6* and *Klhl6* (Fig. 3c and d). The ORR1E elements around these two genes were less accessible in BPA10mg male but more accessible in As female comparing to controls (Fig. 3c), and the expression pattern of these two genes under exposure matched the accessibility changes (Fig. 3d). We next performed transcription factors (TF) binding motif analysis and found that 19 of 20 DA-ORR1Es contained binding motifs of *Ikzf3* and *Elf3*, which are TFs known to regulate immune functions^41–43^ (Fig. 3e and Supplementary file 11). The binding motifs of *Ikzf3* (AGGAAG) and *Elf3* (CTTCCT) were well-conserved in DA-ORR1E and ORR1E consensus sequence, and formed an *Ikzf3*-*Elf3*-*Elf3* pattern within the core region of ORR1E subfamily members (Fig. 3f, S-fig. 4a and Supplementary file 12). The expression levels of DA-ORR1Es associated immune genes were up-regulated in As-exposed female livers and down-regulated in BPA-exposed male livers, agreeing with the accessibility changes of DA-ORR1Es in response to these exposures (S-fig. 4b). We extended motif analysis to all 258 accessible ORR1Es in our data, and multiple sequence alignment indicated 159 accessible ORR1Es preserved at least one of three binding motifs in their core regions (Fig. 3g and Supplementary file 13). Genes around these ORR1Es were enriched in immune-related biological processes and mouse phenotypes (Fig. 3g). Notably, the core regions were more conserved in the 159 ORR1Es with the binding motifs than in the 99 ORR1Es without the motifs (Fig. 3g).

**Figure 3.**
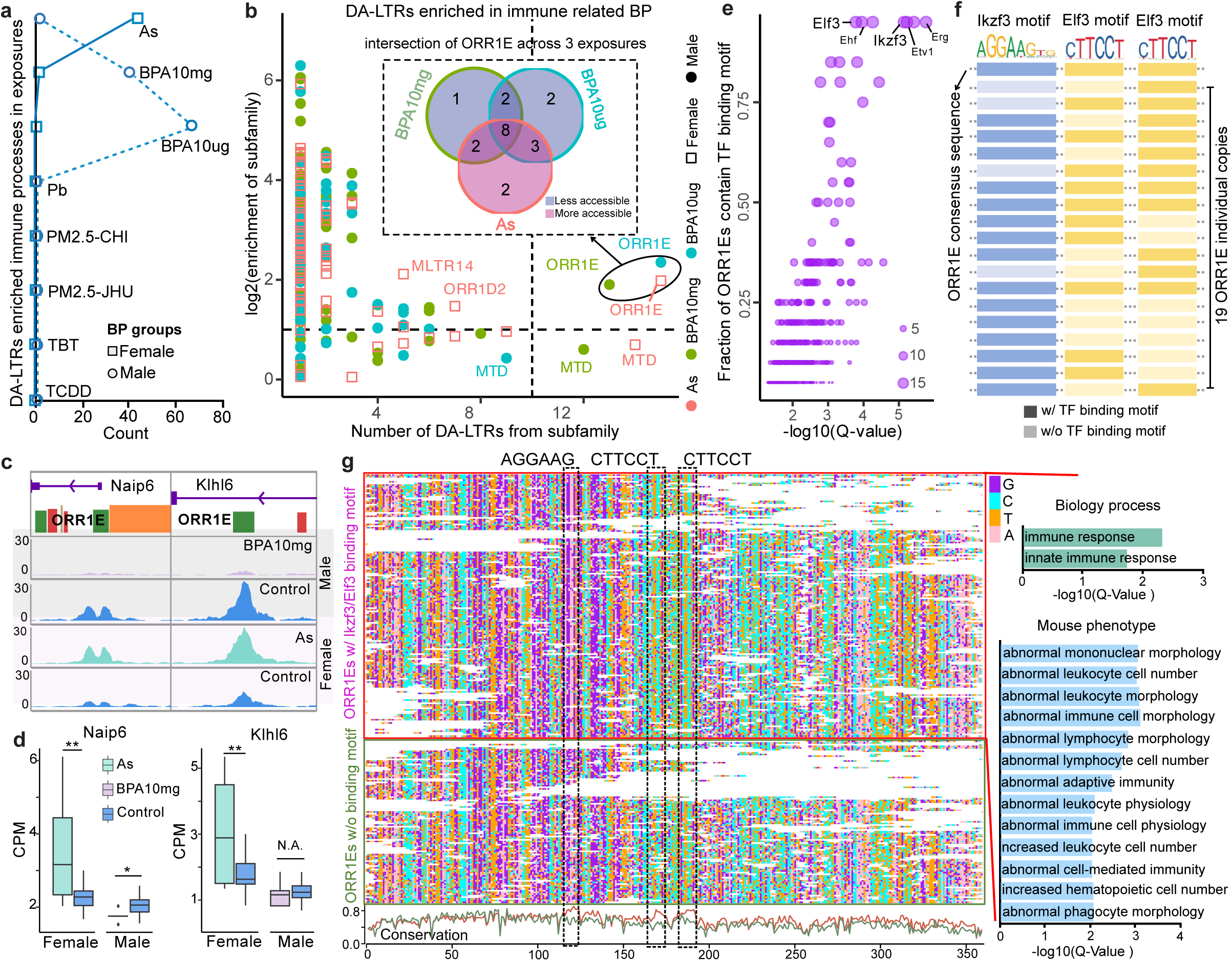
Differentially accessible TEs associated with immune-related processes. **a**) Number of significantly enriched immune-related processes around DA-LTRs (differentially accessible TEs of LTR) in mouse livers with different exposures. Square: Female; Dot: Male; BP: biological process. **b**) Subfamily enrichment of DA-LTRs associated with immune-related processes. X-axis: number of DA-LTRs of enriched subfamily; Y-axis: log2(enrichment) of immune-related DA-LTR subfamily. The ORR1E subfamilies were enriched in As-exposed female liver and BPA-exposed male liver. The Venn diagram (top): the intersection of 20 ORR1Es individual elements across 3 exposures. **c**) ORR1E-derived TSS of *Naip6* and ORR1E-derived enhancer in *Klhl6* intron were closed in BPA10mg-exposed male livers, but became more open in As-exposed female livers. **d**) Expression of *Naip6* and *Klhl6* genes in BPA10mg and As exposures. (Student t-test. *: p-vale < 0.05; **: p-value < 0.01). **e**) Enriched transcription factor binding motifs in differentially accessible ORR1Es (DA-ORR1Es) associated with immune-related processes. **f**) Diagram of *Ikzf3* and *Elf3* binding motifs in DA-ORR1Es. **g**) Multiple sequence alignments of 258 accessible ORR1Es with or without *Ikzf3*/*Elf3* binding motifs. Right: the enriched GO terms of ORR1Es with *Ikzf3*/*Elf3* binding motifs. Bottom: the ORR1Es conservation of the two groups.

### TE-derived transcripts induced by environmental exposures

TEs can initiate the transcription of host genes^1,11,16,17^. To understand how TE-derived genes respond to environmental exposures, we explored the expression of known TE-derived transcripts, and further identified novel TE-derived chimeric transcripts (**Methods**). There were 781 known TE-derived transcripts expressed (TE-transcript, CPM>1) in our data. Their expression profiles were exposure-specific and could easily distinguish the exposed samples from controls in PCA analysis (Fig. 4a). Surprisingly, 540 of the 781 known TE-transcripts were significantly differentially expressed in at least one exposure condition (DE-TE-transcripts, Fig. 4b and Supplementary file 14), and most of these DE-TE-transcripts were up-regulated in exposed animals, especially in female (Fig. 4c). Furthermore, most of DE-TE-transcripts were exposure-specific (S-fig. 5a and b), and very limited shared DE-TE transcripts were identified in both female and male livers in responding to the same exposure (S-fig. 5c).

**Figure 4.**
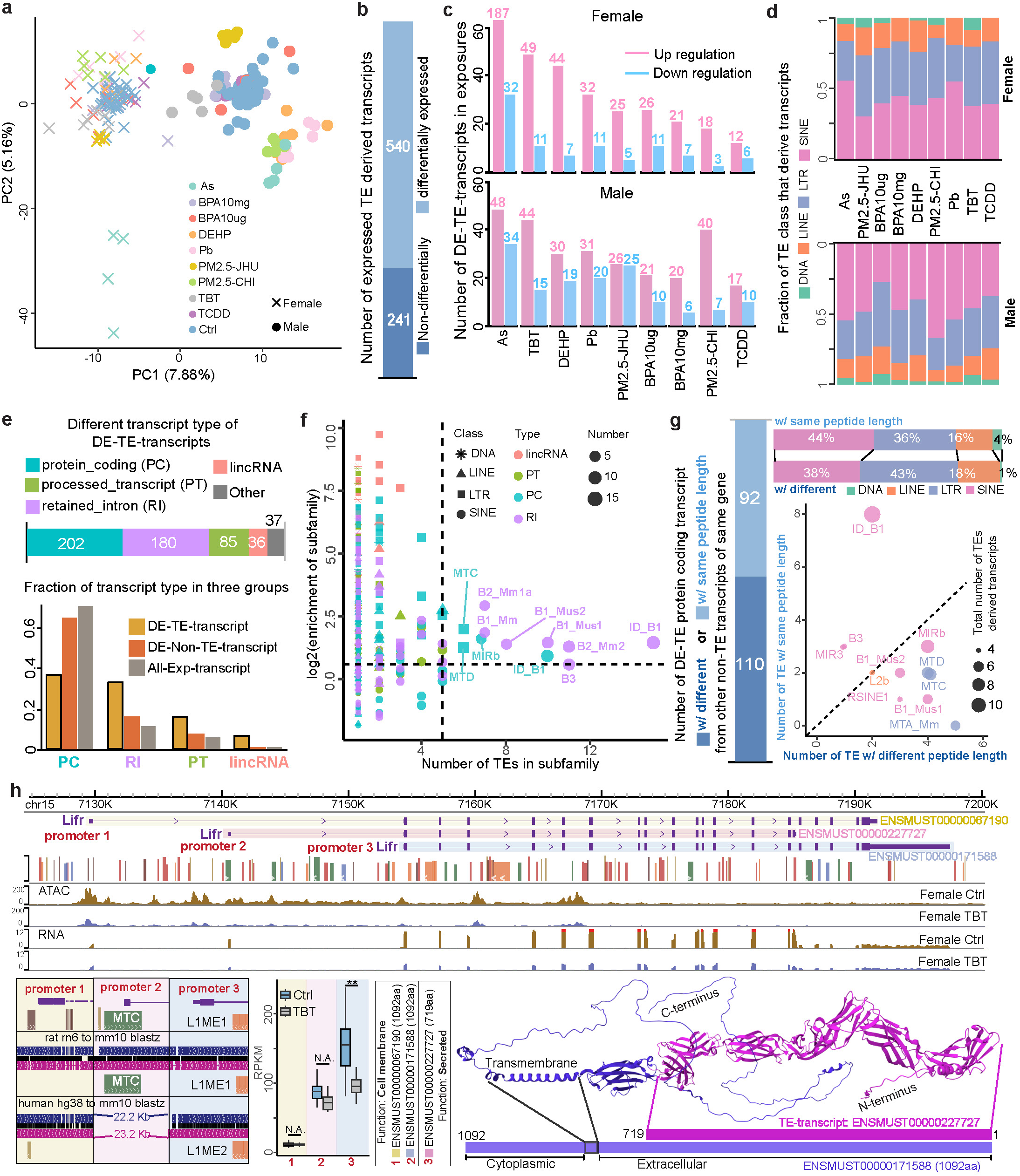
Exposure-specific and sex-specific response of TE-derived transcripts. **a**) PCA plot based on the expression of TE-derived transcripts. **b**) Number of differentially expressed TE-transcripts (DE-TE-transcripts) in response to at least one exposure. **c**) Up- and down-regulated DE-TE-transcripts in female (top) and male (bottom) in response to different exposures. **d**) Class distribution of TEs that derived DE-TE-transcripts in response to exposures. **e**) Transcript-types distribution of DE-TE-transcripts (Top). Higher fractions of retained-intron (RI), processed transcripts (PT), and lincRNA were found in DE-TE-transcripts (Bottom). **f**) Enrichment of TE subfamilies of DE-TE-transcripts. X-axis: the number of TEs in each subfamily. Y-axis: log2 enrichment of TE subfamily in different transcript types. **g**) DE-TE-derived protein-coding transcripts with altered length of protein products (Left), and their TE class and subfamily distribution (Right). **h**) Down-regulation of *Lifr* gene in TBT-exposed females was mainly driven by the loss of expression of MTC-derived transcript isoform (ENSMUST00000227727), which used the intronic promoter (promoter 2) to produce a secreted C-terminal truncated protein (719 AA) that only contains the extracellular domain. (Student t-test. **: p-value < 0.01). Protein folding structure predicted by AlphaFold2.

We noticed near 80% of differentially expressed TE-transcripts used SINE or LTR elements to initiate transcription (Fig. 4d), and various transcript types can be found in our analysis, including 202 protein-coding transcripts, 180 transcripts with retained introns, 85 processed transcripts, 36 lincRNAs, and 37 other types of transcripts (Fig. 4e). Notably, the high proportion of TE-derived retained-intron transcripts in responding to exposure was observed when comparing to non-TE-derived transcripts (Fig. 4e). We next examined the enrichment of specific TE subfamilies across different transcript types among DE-TE-transcripts, and found that seven SINE subfamilies were significantly enriched in transcripts with retained introns, while two LTR and two SINE subfamilies were enriched in protein-coding transcripts (Fig. 4f). Gene Ontology (GO) analysis of DE-TE-transcripts originating from these enriched subfamilies revealed distinct functional associations: retained-intron transcripts were linked to lipid metabolism, DNA repair, and mitochondrial ATP synthesis, whereas protein-coding transcripts were predominantly associated with immune-related processes (Supplementary Fig. 5d).

Among the 202 protein-coding DE-TE-transcripts annotated by Ensembl as capable of translation, 110 were found to produce protein products with altered amino acid sequences compared to their canonical non-TE-derived isoforms, while the remaining 92 TE-transcripts were to produce the same protein products (Fig. 4g and Supplementary file 15). For example, there were two upstream ORR1D1 elements provided start sites for initiating the alternative transcription of the *Hsd3b3* gene and produced the same protein as the non-TE-derived isoform. TE derived isoforms of *Hsd3b3* gene were up-regulated in TBT-exposed male livers but down-regulated in TCDD-exposed female livers. (S-fig. 6). These protein-coding TE-transcripts were mainly derived from LTR, SINE, and LINE elements (Fig. 4g). In particular, MTA_Mm, B1_Mus1, MTC, and MTD elements significantly contributed to these diverse protein products (Fig. 4g). As a type I cytokine receptor, *Lifr* was recently reported as the key factor to stimulate proliferation and differentiation of adipose progenitor cells in internal organs during aging^44^. We discovered that *Lifr* can be transcribed from a murine-specific MTC-derived TSS and two other conserved non-TE-derived TSSs, and produce two distinct protein products (Fig. 4h). The long peptide (1092 aa) from non-TE-derived TSS is a cell surface protein with a transmembrane domain, which is truncated in MTC-derived short peptide (719 aa), and leads *Lifr* to be secreted outside of cells^45–47^. Under early-life exposure to TBT, the male livers significantly down-regulated the MTC-derived transcript but not the canonical isoforms (Fig. 4h).

Previous studies highlighted the importance of reactivated TEs in forming chimeric transcripts, especially in human diseases^1,48,49^. To identify reactivated TEs and their associated chimeric transcripts, we incorporated the TEProf3^50^ method into a customized pipeline to discover TE-derived transcripts in mouse livers exposed to early-life environmental toxicants (**Methods**, S-fig. 7). In total, we discovered 164 novel TE-derived transcripts and classified them in four categories based their relationship with known coding and non-coding genes (Fig. 5a and Supplementary file 16). 89 TE-transcripts were novel TE-chimeric-transcripts of known protein-coding genes, and 20 were TE-chimeric-transcripts of non-coding genes. We also identified 55 novel TE-transcripts, which were not recorded in current gene annotation, including one transcript located within a single TE locus (Fig.5a). Most of these novel TE-transcripts were derived from LTR elements, highlighting the reactivation potential of LTRs (Fig. 5b). We further found 140 of the 164 novel TE-transcripts were significantly affected by different exposures (Fig. 5c and Supplementary file 17), most of which were up-regulated in responding to early-life exposures (Fig. 5d, S-fig. 8 a and b). However, exposures to As, TCDD, DEHP, and Pb resulted in a higher number of downregulated novel TE-transcripts, suggesting exposure-specific regulatory effects on these transcripts (Fig. 5f). Most of these novel TE-transcripts were dysregulated in response to a single exposure, indicating the high exposure-specific sensitivity of TEs to environmental stimulus (S-fig8. c and d). *Cebpe* is a bZIP transcription factor belonging to the CCAAT/enhancer binding protein (C/EBP) family, known to play a critical role in the terminal differentiation and functional maturation of committed granulocytes^51,52^. We identified that *Cebpe* transcription in the liver predominantly arose from a previously unannotated TSS embedded within an RLTR20C2_MM transposable element, located approximately 7.9 kb upstream of the canonical promoter (Fig. 5e). Notably, RLTR20C2_MM-derived *Cebpe* isoform was significantly upregulated in mice exposed to PM2.5-CHI, suggesting that air pollution may alter liver function through TE-mediated transcriptional regulation.

**Figure 5.**
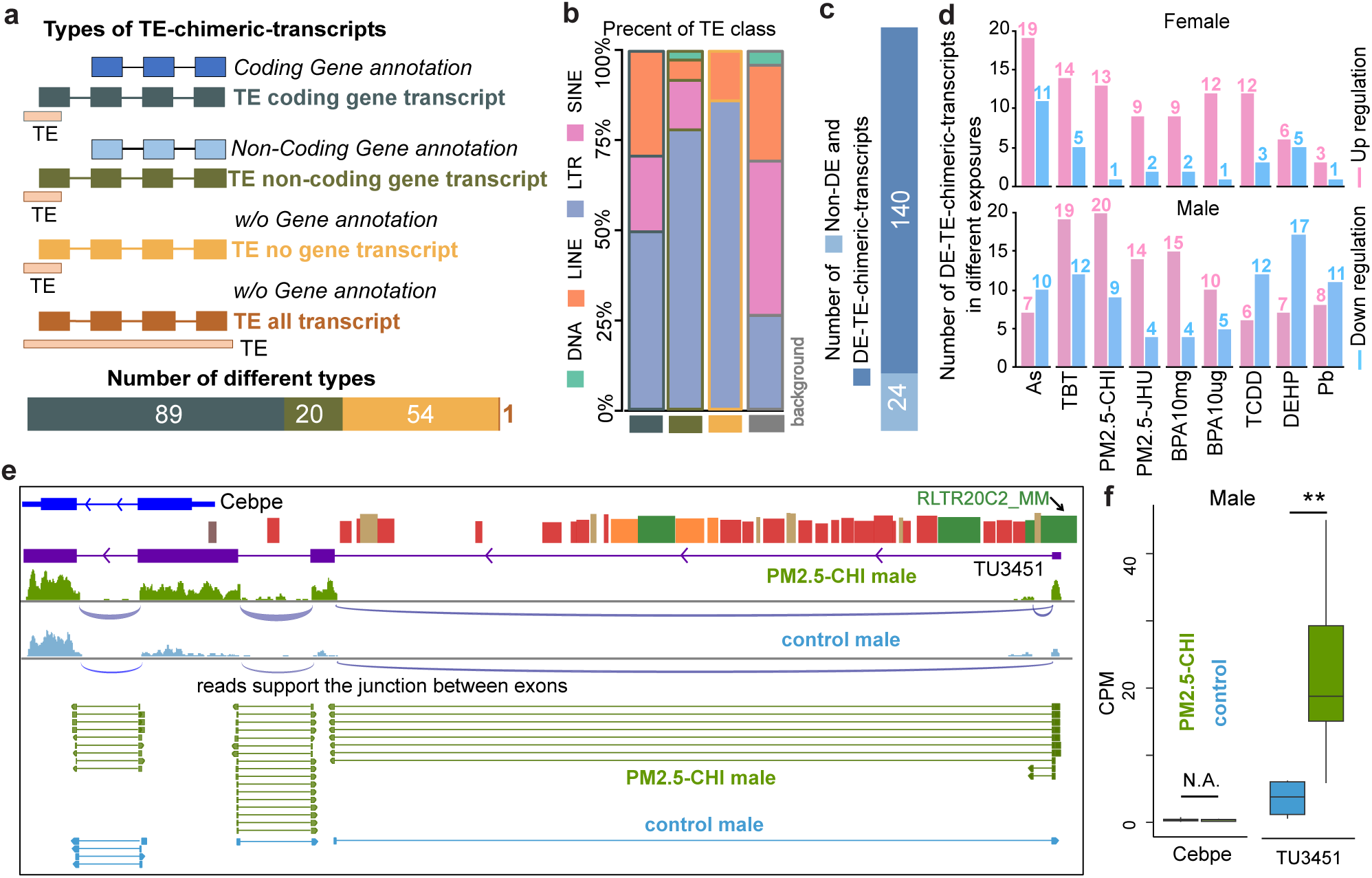
Identification of novel TE-derived transcripts in response to environmental exposures. **a**) Categories of TE-derived chimeric transcripts (TE-chimeric-transcripts) identified in toxicant-exposed livers. TE-derived transcripts can be novel chimeric with annotated coding or non-coding genes, or without known annotation. **b**) Class distribution of TEs that derived novel transcripts. Background fraction indicated the genomic proportion of each TE class in the mouse genome. **c**) Number of differentially expressed novel TE derived transcripts (DE-TE-chimeric-transcripts) in response to at least one exposure. **d**) Up- and down-regulated DE-TE-chimeric-transcripts in response to exposures in female (top) and male (bottom) livers. **e**) Exon-junction analysis of RNA-seq data indicated that RLTR20C2_MM-derived novel chimeric transcript (*TU3451*) of *Cebpe* gene was the dominant transcript isoform in male liver. **f**) Expression of canonical *Cebpe* and RLTR20C2_MM-derived novel chimeric transcript (*TU3451*) in male PM2.5-CHI-exposed liver and control livers (student t-test. **: p-value<0.01).

In TBT-exposed female livers, we discovered that an RLTR34C_MM element located 3.63kb upstream of the *Pfas* gene, which is involved in the purine biosynthetic pathway^53^, can derive a novel chimeric transcript (*TU1781*) of *Pfas* (Fig. 6a). This novel RLTR34C_MM-*Pfas* transcript also included three novel exons during transcription and splicing. ORF analysis indicated that *TU1781* added an extra 195 bp coding region to *Pfas*, which could result in an extended N-terminus (65 AA) at the protein level (Fig. 6b-c). Transcript-level expression analysis revealed a significant isoform switch between the canonical *Pfas* and the RLTR34C_MM-*Pfas* chimeric isoform. While the canonical *Pfas* isoform was predominantly expressed in control livers, TBT exposure reactivated the RLTR34C_MM element, leading to the dominant expression of the novel chimeric transcript (Fig. 6b). We also identified 55 novel TE derived transcripts that did not splice together with any known coding or non-coding genes. For example, an RMER15 elements initiated two chimeric transcripts (*TU7311* and *TU7312*) which were located in the intron of the *Cfi* gene in an antisense orientation (Fig. 6d and S-fig. 9). RNA sequencing reads supporting the junctions between the exons of these two chimeric transcripts and one of the transcripts was predicted to generate a peptide sequence with helix protein structures (Fig. 6e). These two chimeric transcripts were highly expressed in As males, but the *Cfi* gene showed low expression in As male (Fig. 6f).

**Figure 6.**
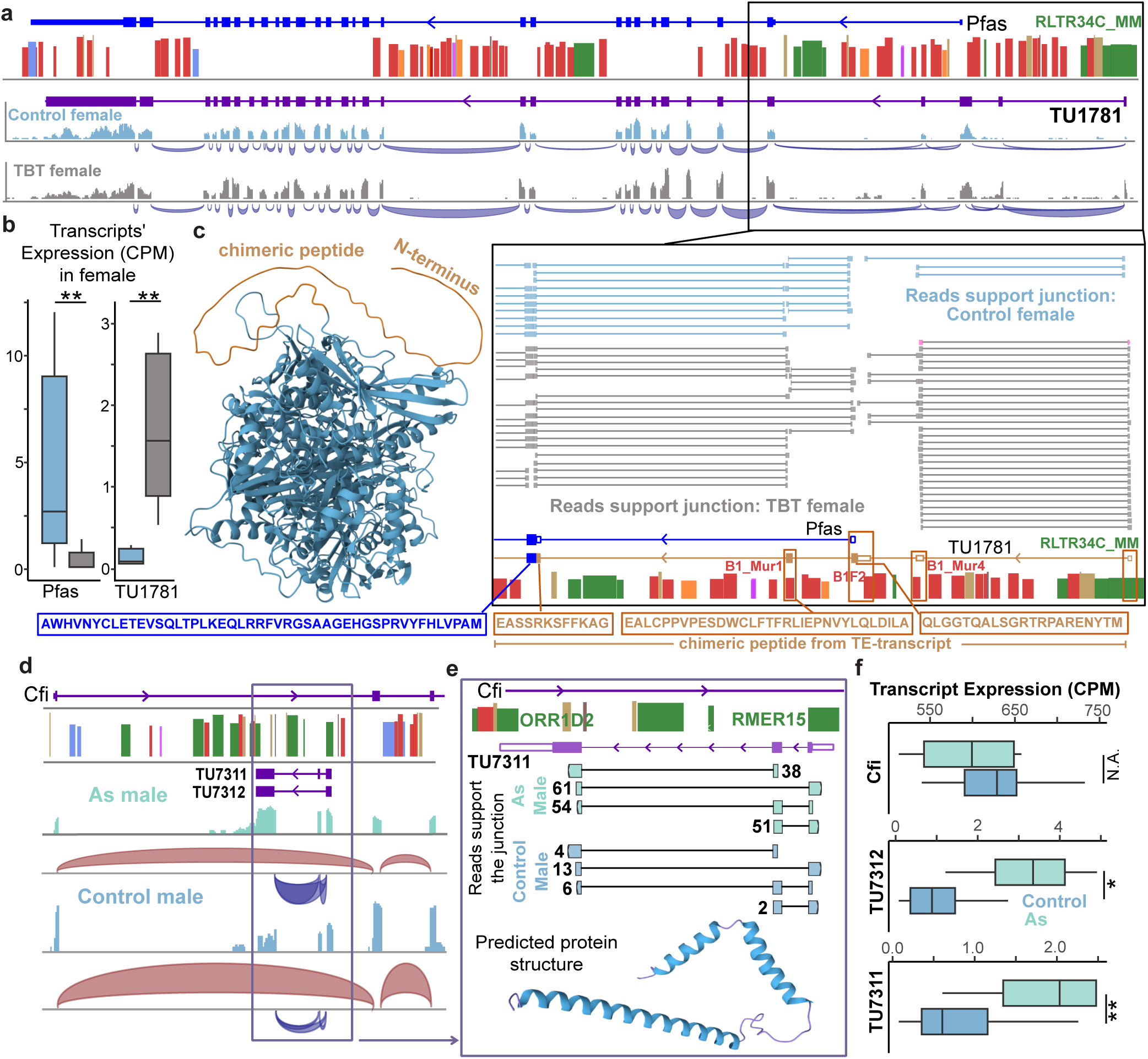
TEs can derive novel chimeric transcripts and unannotated genes. **a**) Exon-junction analysis of RNA-seq data indicated RLTR34C_MM-derived novel chimeric transcript of *Pfas* gene in TBT-exposed female liver. Exon-junction analysis (Bottom) indicated RLTR34C_MM-derived 1^st^ exon can be spliced with the three downstream exonized Alu elements, which can provide novel peptide coding capacity. **b**) Anti-correlated expression changing pattern of canonical *Pfas* and RLTR34C_MM-derived novel chimeric transcript (*TU1781*) (student t-test. **: p-value<0.01). **c**) Predicted protein structure (AlphaFold2) of *TU1781* contained an uncharacterized N-terminal peptide sequence (65 AA). **d**) RMER15-derived novel transcripts located in the 1^st^ intron of *Cfi* gene and exhibited antisense orientation of *Cfi* gene in male liver. **e**) Exon-junction analysis indicated alternative splicing patterns between RMER15-derived 1^st^ exon and other exons. *TU7311* transcript contains a 4209 bp ORF, which can be translated to 1402 AA peptide (with protein structure predicted by AlphaFold2). **f**) Two TE-derived transcripts (*TU7311*, *TU7312)* were significantly up-regulated in As-exposed livers, while expression of *Cfi* was relatively stable. (*: p-value<0.05; **: p-value<0.01).

## Discussion

Transposable elements (TEs) constitute a substantial portion of mammalian genomes and contribute to gene regulation, chromatin organization, and the modulation of host genomes in response to external stimuli^1–3,5,6,8^. However, the roles of TEs in response to different environmental toxicants remain largely unknown. Here, we conducted a comprehensive analysis to explore the regulatory roles of TEs in adult mouse liver and how these TEs respond to various early-life environmental exposures, including As, Pb, BPA, TBT, DEHP, TCDD, and PM2.5. Our integrative analysis of the transcriptome and epigenome revealed sustained and dynamic activity of TEs in adult livers five months after early-life exposures, emphasizing TEs as critical regulatory elements in mediating long-term responses to environmental stress.

We found that approximately 20% of open chromatin regions were derived from TEs. 6,699 TEs displayed significant differences in accessibility as a function of environmental exposures. These TEs were mainly located in intron and intergenic regions with potential cis-regulatory functions, underscoring the impact of TEs on chromatin landscapes in response to external stimuli. Our results are consistent with previous studies demonstrating TEs’ role as functional regulatory elements influencing gene expression. Furthermore, GO analysis revealed that differentially accessible LINEs were enriched for metabolic-related processes, and LTRs were enriched for immune responses. In particular, the ORR1Es subfamily was highly associated with immune-related transcription factor binding motifs, suggesting that exposures could induce different accessibility patterns of specific TEs, which then mediate distinct biological functions.

Previous studies indicated that TEs can initiate gene transcription to increase transcripts and protein diversity, and could contribute to the adaptability of hosts in various environmental stresses^3–6,16,17,19^. Our transcriptomic analysis revealed that a substantial proportion (∼70%) of the 781 identified TE-derived annotated transcripts were differentially expressed in response to environmental exposures, with a majority being up-regulated. In addition, we discovered 164 novel TE-chimeric transcripts, ∼85% of which were significantly affected by environmental exposures. Both results emphasized the high sensitivity of TEs in response to toxicants exposure. These TE-derived transcripts also represented an evolutionary mechanism of gene regulatory network to adapt the continuously changing environmental stress. For example, one murinae-specific RLTR34C_MM was adopted to initiate a novel transcript isoform of the *Pfas* gene in response to TBT exposure (Fig. 6c). This novel chimeric transcript also contained three exonized SINE elements, resulting in an extended N-terminus (65 AA) of the canonical *PFAS* protein. The functional consequence of such novel proteins remained to be investigated, but our current results highlighted the impact of TEs on transcriptional innovation and contributing to environmental adaptation.

The sex-specific response of TEs to those environmental exposures was observed at both chromatin accessibility and expression levels in all exposed conditions, suggesting that sex-specific epigenetic regulation and gene expression of animals when responding to environmental stress. This sex-specific pattern of TEs indicated the importance of understanding the molecular mechanisms behind sex-biased diseases, especially those associated with environmental exposures, such as metabolic and immune disorders. We also observed that a small fraction of TEs were simultaneously affected by different environmental exposures, suggesting converged pathways involving some TEs in response to different environmental stimuli.

While our study uncovered these novel insights, it is still subject to certain limitations. First, we focused on adult liver, which cannot fully capture the systemic impact of environmental exposures on TE activity across diverse tissues and developmental stages. Second, bulk tissue profiling limited our ability to resolve cell-type-specific effects, and further single-cell approaches could provide a finer-resolution of TE dynamics in response to environmental exposures. Finally, the functional roles of many TE-derived transcripts, particularly the novel non-coding and chimeric transcripts identified in our study, remained largely unexplored. Although a significant proportion of these transcripts were differentially expressed in response to environmental exposures, further functional validation would provide more clues to their biological relevance.

In conclusion, we described the full landscape of TE activation in adult livers, and characterized their long-term chromatin accessibility modulation and TE-derived gene expression in response to early-life toxicant exposures. The dynamic regulation of TEs, particularly those from younger TE subfamilies, revealed the importance of TE-derived regulatory elements in environmental adaptation during animal evolution. These findings offer new insights into the molecular mechanisms by which environmental stressors influence genome regulation and underscore the potential of transposable elements as biomarkers or therapeutic targets in the context of environmental health.

## Method

### Animal exposure paradigm and tissue collection

C57BL/6 (B6; Jackson Laboratory, Bar Harbor, ME) mice were used for all experiments, except for the lead (Pb) and phthalate (DEHP) exposure studies, which employed wild-type non-agouti (a/a) mice derived from a >230-generation colony of viable yellow agouti (Avy) mice.

Mice were exposed to toxicants perinatally via maternal diet, drinking water, or air breathing, with the exposure window spanning from pre-conception through weaning. In brief, two weeks before mating, virgin female dams (6–8 weeks old) were randomly assigned to the exposure or control group.

Detailed methods for the animal exposure and tissue collection have been previously described in our flagship paper.

### Transposable element annotation data

The transposable elements (TEs) data of mouse (mm10) was obtained from the UCSC database (http://hgdownload.soe.ucsc.edu/goldenPath/mm10/database/)^54^. Simple repeats and satellites were not included in this TE dataset, which contained TEs of DNA, LINE, SINE and LTR classes. The evolutionary conservation information of TE subfamily in mouse was gathered from the Dfam database (https://www.dfam.org/releases/Dfam_3.8/)^55^. Then, taxa information of TE subfamily was extracted and the subfamilies from the Mus_genus and Mus_musculus were assigned to Mus_musculus clade, TE subfamilies from Muridae and Murinae belonged to Muridae clade, subfamilies of Glires and Rodentia were assigned to Rodentia.

### Raw sequence data and processing

Raw fastq files of RNA-seq and ATAC-seq data were gathered from liver samples of 5 months mice exposed to environmental toxicants during early development, including arsenic (As), lead (Pb), bisphenol A (BPA), tributyltin (TBT), di-2-ethylhexyl phthalate (DEHP), 2,3,7,8-tetrachlorodibenzo-p-dioxin (TCDD), and particulate matter less than 2.5 µm in diameter (PM2.5).

Raw fastq files of RNA-seq data from different exposures were processed by Cutadapt (v1.16; --quality-cutoff=15,10 --minimum-length=36), FastQC (v0.11.7), and STAR (v2.5.4b; --quantMode TranscriptomeSAM --outWigType bedGraph --outWigNorm RPM) for trimming, QC report and mouse genome mapping (mm10) with our own built pipeline, the TaRGET-II-RNA-seq-pipeline (https://github.com/Zhang-lab/TaRGET-II-RNAseq-pipeline)^56–58^. Then, the genes and transcripts’ expressions across normal and exposed samples were separately calculated by featureCounts (v1.5.1) and RSEM (v1.3.0) within the pipeline based on GENCODE vM20 gene annotation of mouse genome^59–61^.

Raw fastq of ATAC-seq data were processed by an in-house pipeline, the TaRGET-II-ATAC-seq-pipeline (https://github.com/Zhang-lab/TaRGET-II-ATACseq-pipeline) integrated AIAP packages contained optimized QC report and analysis pipeline with default parameters to generate the open chromatin regions (OCRs)^62^. Then, consensus regions of OCRs across different exposure and control samples were generated with Index (https://github.com/Altius/Index) method used for downstream analysis^63^. The ATAC-seq signals of consensus OCR were calculated by using the intersectBed method of bedtools (v2.25.0)^64^.

The TaRGET-II-WGBS-pipeline (https://github.com/Zhang-lab/WGBS_analysis) was built by using the Cutadapt (v1.16; --quality-cutoff=15,10 --minimum-length=36) and Bismark (v0.19.0; --bowtie2 -X 1000 --score_min L,0,-0.6 -N 0 --multicore 2 -p 4) to do the trimming and mouse genome mapping^65^. The pipeline also incorporated quality control, generated user-friendly files for computational analysis and output genome browser tracks for data visualization. Next, the methylation status for each aligned CpG was calculated using Bismark Methylation Extractor (v. 0.7.10) at a minimum of 20X coverage per site in a strand-specific manner (run-time parameters: -p, no_overlap, --comprehensive, --bedGraph)^65^. Then, the averaged methylation of TEs were calculated based on the CpG within the TEs.

The RNA-seq and ATAC-seq datasets were normalized to correct for batch effects arising from different data production centers by using the method described in the flagship paper, and then the consortium-normalized read count data were used for downstream analysis. The differentially expressed transcripts were identified by edgeR with FDR < 0.01 and abs(log2(Foldchange)) > 1^66^. The edgeR was used to identify the differential accessible regions (DARs) with FDR < 0.01 and abs(log2(Foldchange)) > log2(1.5).

### The accessibility and expression of TEs at subfamily level

The bam files of RNA-seq and ATAC-seq data were processed by SQuIRE (v0.9.9) and T3E (v1.1) methods, which were separately used to calculate the expression and ATAC-seq signals of TE subfamily^67,68^. The principal component analysis was generated by prcomp function of R package based on expression and ATAC-seq signals of TE subfamilies^69^. The edgeR method was applied to calculate the differentially expressed TE subfamilies (DE-subfamilies) between exposure and control samples with FDR < 0.001 and abs(log2(Foldchange)) > 0.8, and the median CPM of TE subfamily should be greater than 100 in control or exposure samples. The differential accessible TE subfamilies (DA-subfamilies) were identified by edgeR method between exposure and control samples with FDR < 0.001 and abs(log2(Foldchange)) > 0.15, and the median CPM of TE subfamily should be greater than 100 in control or exposure samples. The correlation coefficient between female and male responding to each exposure was calculated by cor function of the R package separately based on the log2(foldchange) of DE-subfamilies and DA-subfamilies. The enrichment of TE class for DA-subfamilies shared between female and male under the PM2.5-CHI was calculated based on the fraction of one class in shared DA-subfamilies divided by the fraction of same class in all TE subfamilies.

### The differentially accessible TEs in different exposures

The OCRs and DARs derived from TEs were defined as summit of OCRs and DARs intersected with TEs. Then, the differentially accessible TEs (DA-TEs) were identified by TEs that intersected with summit of DARs. The ratio, representing the enrichment of subfamilies, was calculated from the fraction of one subfamily in DA-TEs divided by fraction of same subfamily in mouse genome. The intersectBed method of bedtools was used to determine the number of DA-TEs in different genomic features, including transcription start site (TSS), transcription terminate site (TTS), exons, introns, and intergenic regions, which were defined by using the GENCODE vM20 gene annotation of the mouse genome. And the mouse cis-regulation elements (cCREs), including CTCF-only, DNase-H3K4me3, promoter (PLS), proximal enhancer (pELS), and distal enhancer (dELS), were collected from ENCODE database used to explore the potential regulatory function of those DA-TEs by intersectBed method^70^. The enriched biological processes and mouse phenotypes in different classes of those DA-TEs from different exposures were identified by GREAT (version 4.0.0) with the following analysis settings: (1) Species assembly: mouse, NCBI build 38; (2) Background regions: whole genome; (3) Association rule: basal plus extension. (4) filtered with cutoffs of Binom FDR Q-Val < 0.05^71^. The enrichment analysis of TE subfamily was measured from fraction of one subfamily in DA-TEs associated with metabolic or immune processes divided by fraction of the same subfamily in mouse genome. The transcription factor binding motifs (TFBS) in ORR1Es were identified by FIMO software (4.10.2) based on motif weigh matrix file from JASPAR (JASPAR2022_CORE_vertebrates_non-redundant_pfms_meme)^72,73^. The muscle method (v3.8.31) was used to generate multiple sequence alignment for individual copies of ORR1E and consensus sequence^74^. WashU epigenome browser was used to visualize the RNA-seq and ATAC-seq tracks of examples^75^.

### TE derived transcripts in different exposures

The known transcripts in mouse genome (mm10) were obtained from the GENCODE vM20 gene annotation file. Transcript types, including protein-coding, processed transcripts, retained introns, and lincRNAs, were annotated according to GENCODE classifications. TE derived known transcript (TE-transcripts) was defined as TE intersected with TSS of known transcripts from GENCODE vM20 gene annotation. The principal component analysis was generated by prcomp function of R package based on expression of TE-transcripts. The edgeR method was used to identify the differentially expressed TE-transcripts (DE-TE-transcripts) with the following criteria: FDR < 0.01 and abs(log2(Foldchange)) > 1. The enrichment of TE subfamilies in different types was calculated from fraction of subfamily in different transcript types divided by fraction of the same subfamily in mouse genome. Then, the DAVID method was used to identify biological processes for enriched subfamilies derived protein-coding and retained-intron transcripts^76^. Next, the translated sequences of protein-coding transcripts of vM20 downloaded from GENCODE database were used to identify DE-TE-protein-coding transcript with different or same peptide length from other non-TE transcripts of the same gene. The WashU epigenome browser was used to show the example of TE derived known transcript of *Lifr* gene. And the protein structure was gathered from AlphaFold Protein Structure Database^77,78^.

To identify TE derived chimeric transcripts (TE-chimeric-transcript) induced by different exposures, fastq files of RNA-seq samples were re-processed by STAR to generate bam files considering multiple alignment of reads with following parameter, --outFilterMultimapNmax500 and --outSAMattributesNH HINM MD XSAS. Then, these bam files were processed by TEProf3^50^ (TE-derived Promoter Finder 3, https://github.com/Yonghao-Holden/TEProf3) to identify TE-derived promoters and transcripts. Next, TE-derived transcripts that TE derived first exons intersected with known exons from GENCODE vM20 annotation were filtered in the analysis. The junctions between 1^st^ and 2^nd^ exon were supported by at least 2 reads in individual sample and transcript’s CPM ≥ 1 in more than half of samples from one condition. The TE derived chimeric transcripts were classified into 4 different groups by interacting with annotated coding and non-coding genes from GENCODE vM20. TE coding and non-coding gene transcript was TE-chimeric-transcript that separately overlapped with annotated coding and non-coding gene. Among the transcripts without intersection with annotated genes, if the whole transcript came from TE, it was defined as TE transcript, otherwise the transcript belonged to TE no gene transcript. The differentially expressed TE-chimeric-transcripts (DE-TE-chimeric-transcripts) were identified by edgeR with the FDR < 0.05 and abs(log2(Foldchange)) > log2(1.5).

## Data availability

RNA-seq, ATAC-seq and WGBS data in the paper has been deposited through Gene Expression Omnibus (GEO) repository: GSE146508 and those data are also available at the TaRGET II data portal (https://data.targetepigenomics.org/).

TaRGET-II-RNA-seq-pipeline: https://github.com/Zhang-lab/TaRGET-II-RNAseq-pipeline

TaRGET-II-ATAC-seq-pipeline: https://github.com/Zhang-lab/TaRGET-II-ATACseq-pipeline

TaRGET-II-WGBS-pipeline: https://github.com/Zhang-lab/WGBS_analysis

Custom codes are available at GitHub: https://github.com/BenpengMiao/TaRGET-TE

## Authors’ contributions

BZ and TW conceived and designed the study. MB and CW supervised the study. BM performed the computational analysis and data visualization. YZ, WS, and SF processed the raw sequence data. BM, BZ, and TW wrote the manuscript.

## Funding

This work was supported by National Institute Health: U24ES026699 and R35GM142917.

## Competing interests

The authors have declared no competing interests.

## Supplementary files

Supplementary file 1. Sample summary of RNA-seq and ATAC-seq data including exposure, sex, tissue, and stage information.

Supplementary file 2. Differentially expressed TE subfamilies responding to different exposures in 5 months mouse liver.

Supplementary file 3. Differentially accessible TE subfamilies responding to different exposures in 5 months mouse liver.

Supplementary file 4. Correlation between female and male in response to exposures separately based on the expression and accessibility of TE subfamilies.

Supplementary file 5. Class enrichment of differentially accessible TE subfamilies shared between female and male in PM2.5-CHI exposure.

Supplementary file 6. The chromatin accessibility and methylation level of TE subfamilies from LINE shared between female and male in PM2.5-CHI.

Supplementary file 7. The differential accessible TEs (DA-TEs) induced by exposures in female and male of 5 months mouse liver.

Supplementary file 8. The subfamily enrichment of differential accessible TEs under different environmental exposures in female and male of 5 months mouse liver.

Supplementary file 9. Enriched biology processes based on different classes of less and more accessible DA-TEs in different exposures of 5 months mouse liver.

Supplementary file 10. Differential accessible TEs of LTR associated with immune process under environmental exposures.

Supplementary file 11. The transcription factor binding motifs in differential accessible ORR1Es associated with immune process.

Supplementary file 12. The multiple sequence alignment of ORR1E consensus sequence and differential accessible ORR1Es associated with immune under exposure.

Supplementary file 13. The multiple sequence alignment of accessible ORR1Es.

Supplementary file 14. Exposures induced differentially expressed known TE-derived transcripts.

Supplementary file 15. TE-derived protein coding transcripts with different peptide from non-TE derived transcripts of same gene.

Supplementary file 16. The TEProf3ut output of TE-derived chimeric transcripts.

Supplementary file 17. Differentially expressed TE-derived chimeric transcripts responding to environmental exposures.

