## supplementary figures for "The impact of environmental exposures on the epigenomic and transcriptomic landscape of transposable elements"

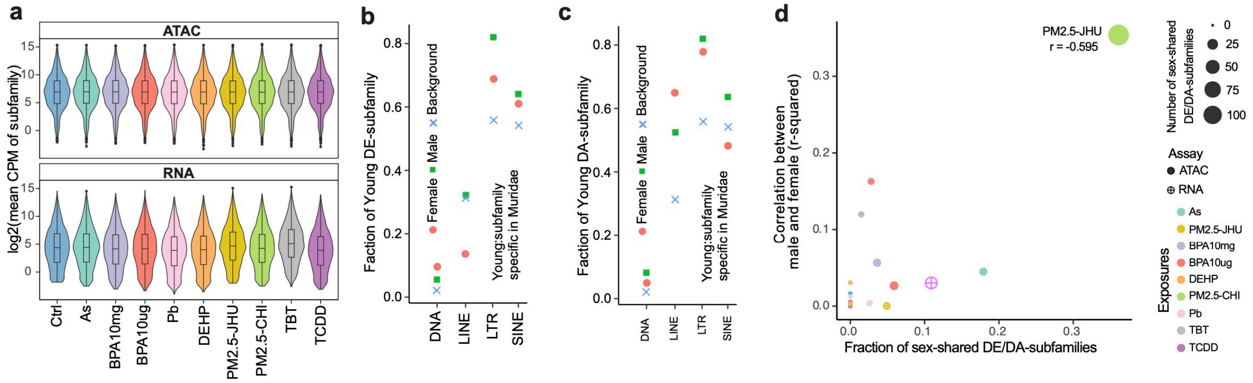


**S-figure 1.** Exposures induced expression and chromatin accessibility changes of transposable elements (TE) at subfamily level. **a**) Boxplot showed averaged ATAC-seq signal and expression value of TE at subfamily level across control and exposure samples. The expression and accessibility of TE subfamilies responding to different exposures were comparable to the control samples at subfamily level. **b**) The faction of DE-subfamilies from exposures belonged to young subfamily separately for DNA, LINE, LTR and SINE. The Muridae specific subfamily was defined as young subfamily. The background is fraction of total subfamilies from DNA/LINE/LTR/SINE in mouse genome belonged to young group. The DE-subfamilies from LTR and SINE showed higher fraction of young group than the background. **c**) The faction of DA-subfamilies from exposures belonged to the young subfamily separately for DNA, LINE, LTR and SINE. The DA-subfamilies from LTR and LINE displayed higher fraction of young group than the background. **d**) Correlation between female and male responding to different exposures separately for RNA-seq and ATAC-seq samples. The y axis was value of the correlation coefficient calculated separately based on the expression and open signals of TE subfamilies for RNA-seq and ATAC-seq data, the x axis was the fraction of DE/DA-subfamilies shared between female and male in different exposures. PM2.5-CHI induced high negative correlation and high fraction of shared DA-subfamilies between female and male. Shapes: assay; color: different exposures; size: number of DA/DE-subfamilies shared between female and male.

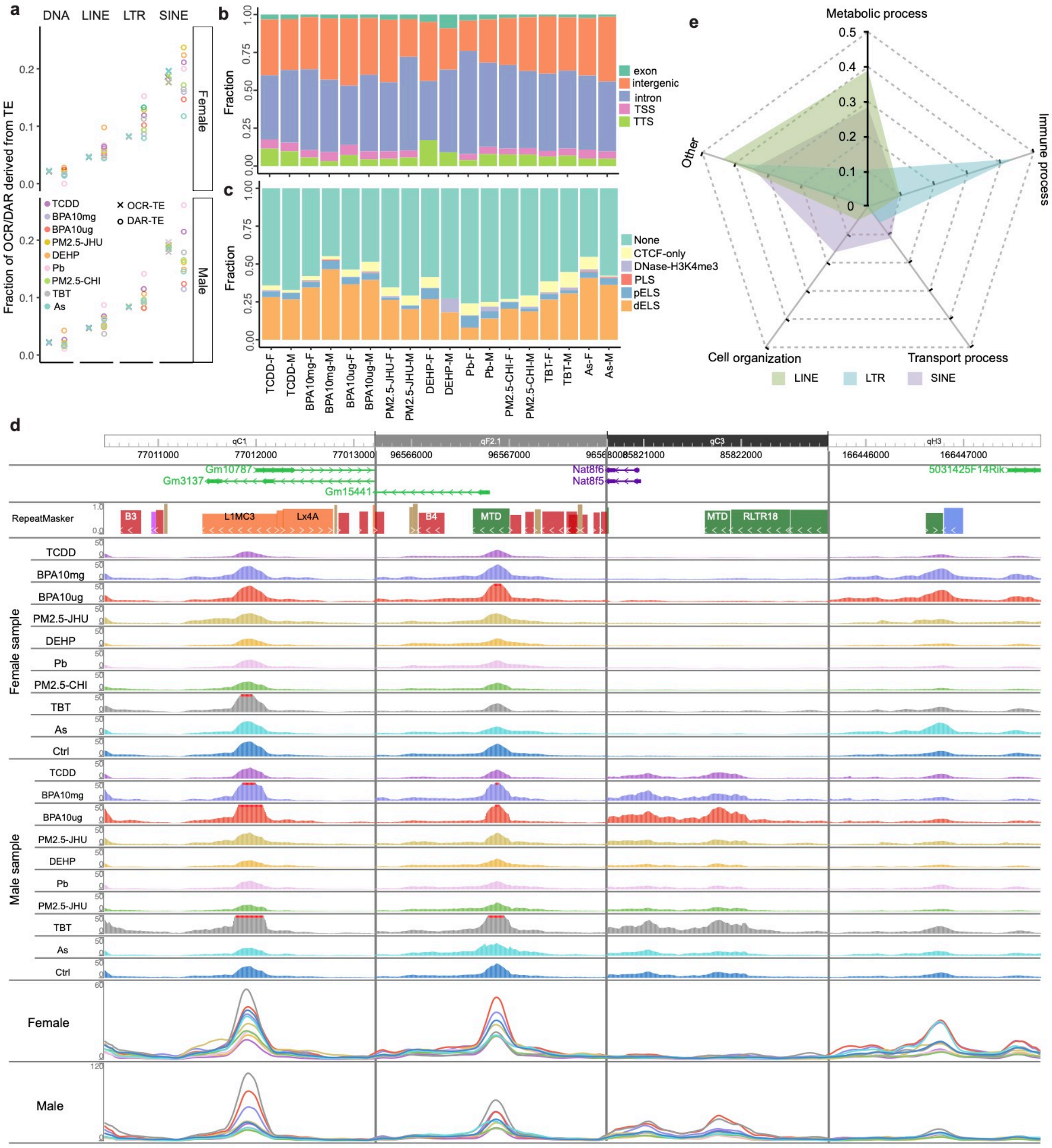


**S-figure 2.** Exposures induced chromatin accessibility changes of individual transposable elements. **a**) Fraction of open chromatin regions (OCRs) and differential accessible regions (DARs) derived from TEs of different TE classes across exposures. The exposures resulted in more dynamic fractions of DAR derived from TEs for both female and male. Dot: DAR derived from TEs (DAR-TE); Cross: OCR derived from TEs (OCR-TE); Colors: different exposures. **b**) Genomic distribution of differential accessible TEs (DA-TEs) in different exposures separately for female and male. Most of DA-TEs located in intron and intergenic regions. **c**) Fraction of DA-TEs annotated as cis-regulatory elements (cCREs) in each exposure separately for female and male, including CTCF-only, DNase-H3K4me3, promoter (PLS), proximal and distal enhancer (pELS and dELS). **d**) Example of DA-TEs showing dynamic responding to exposures. Different colors represented different exposures. **e**) Different class of DA-TEs enriched biology processes under exposures, including metabolic, immune, transport process and cell organization. High fraction of enriched biology processes in LTR associated with immune response, LINE and SINE showed high fraction linked to metabolic process.

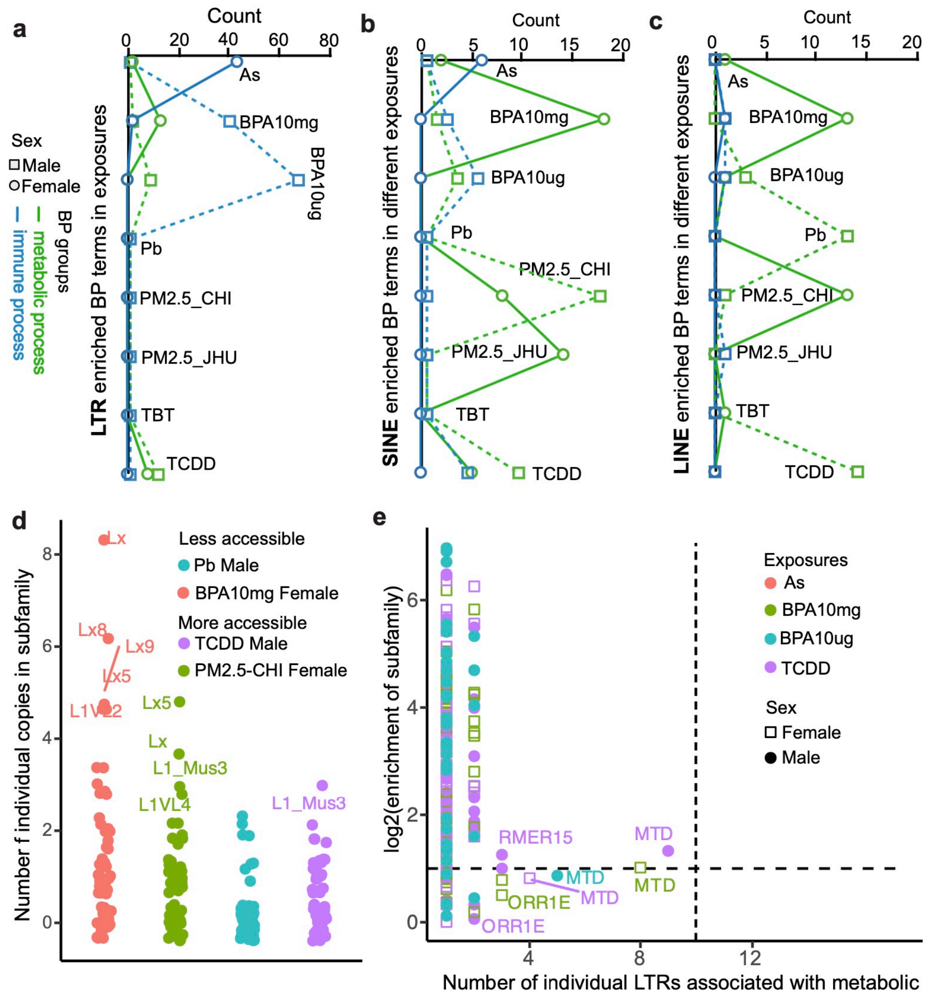


**S-figure 3.** Enriched biology processes identified in different classes of DA-TEs across exposures. Number of biology processes associated with immune and metabolic enriched in **a**) LTR, **b**) SINE and **c**) LINE. Most DA-TEs of LTR enriched biology processes mainly associated with immune. Many SINE and LINE enriched biology processes mainly associated with metabolic process. The number of enriched biology processes were dynamic across different exposures and sex. Square: Male; Dot: female. Green: metabolic process; Blue: immune process. **d**) Number of LINE DA-TEs from different subfamilies associated with metabolic process in male Pb and TCDD, female BPA10mg and PM2.5-CHI. Colors: different exposures. e) Subfamily enrichment of LTR DA-TEs associated with metabolic processes. The x axis was number of DA-TEs from one subfamily associated with metabolic process, y axis was log2 enrichment that was measured from fraction of one subfamily in DA-TEs associated metabolic process divided by fraction of same subfamily in mouse genome. Square: Male; Dot: female.

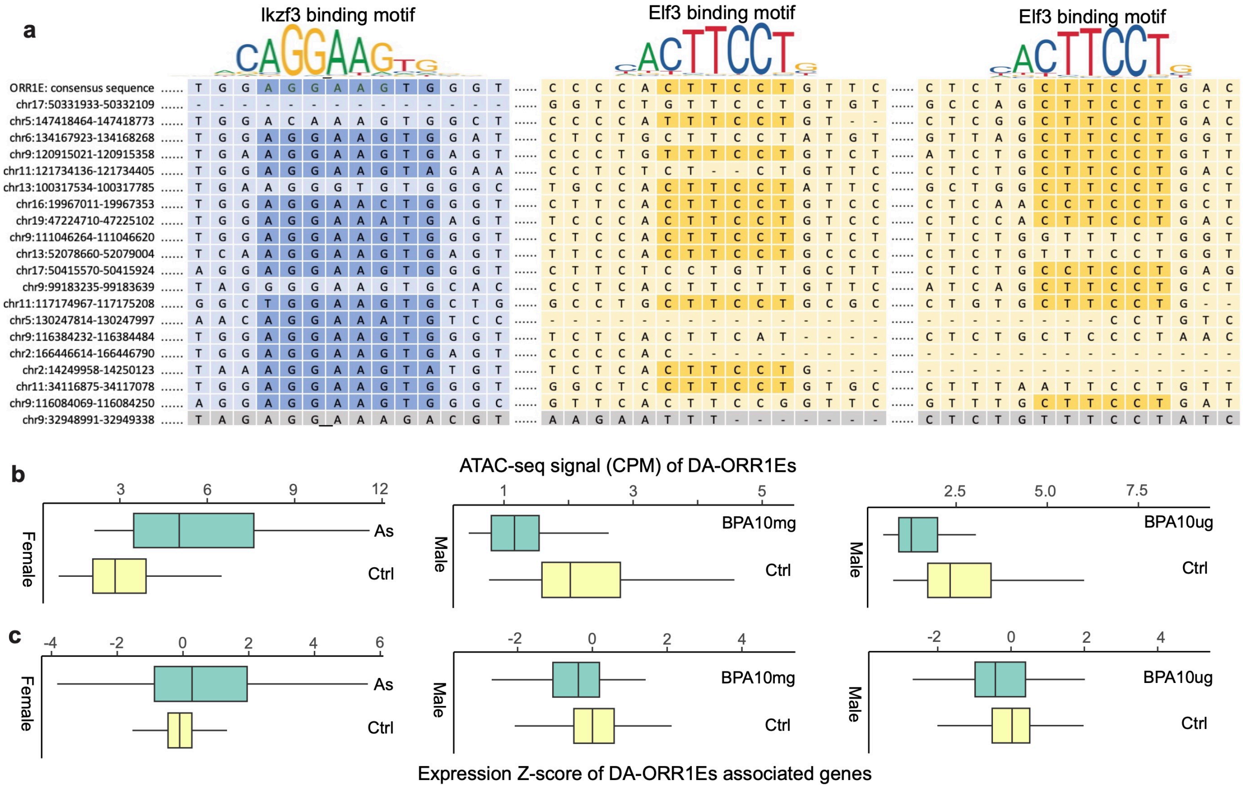


**S-figure 4.** Transcription factor binding motifs within differential accessible ORR1Es (DA-ORR1Es) associated with immune. **a**) The multiple sequence alignments across ORR1E consensus sequence and sequences of 20 DA-ORR1Es. Three core regions with Ikzf3 (AGGAAG) and Elf3 (CTTCCT) binding motifs were identified across those DA-ORR1Es. **b**) ATAC-seq signals of DA-ORR1Es associated with immune in different exposures. DA-ORR1Es identified in As female were more accessible comparing to control. DA-ORR1Es identified in two male BPA exposures were both less accessible comparing to control. **c**) Z-score of expression for DA-ORR1Es linked genes in female As, male BPA10mg and BPA10ug samples. More accessible DA-ORR1Es associated genes showed higher expression in As female than control samples. Less accessible DA-ORR1Es associated genes showed lower expression in both BPA10mg and BPA10ug male than control samples. Green box: exposure samples; Yellow box: control samples.

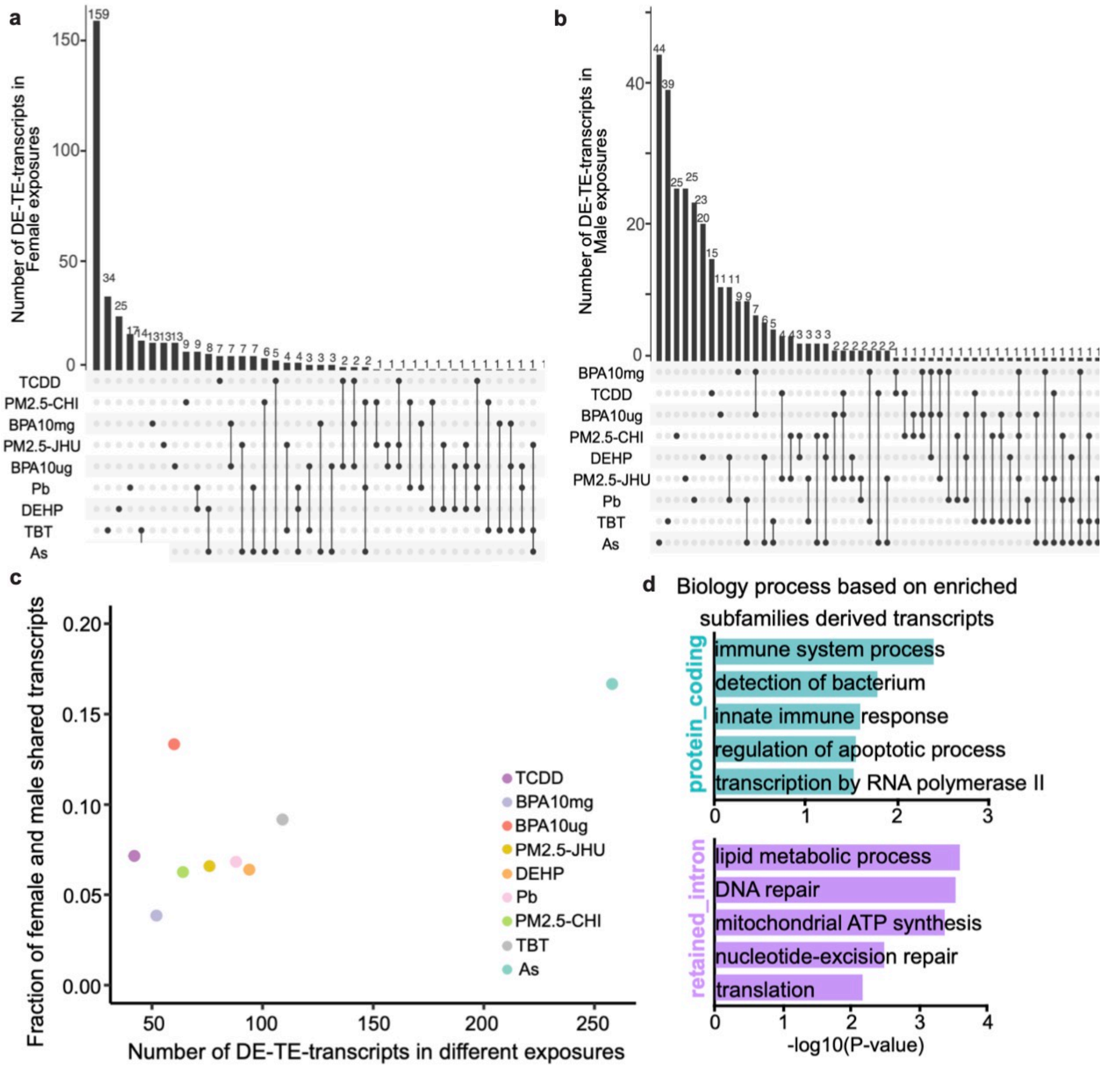


**S-figure 5.** Exposures induced differentially expressed TE derived known transcripts (DE-TE-transcripts). Distribution of DE-TE-transcripts across different exposures in female **a**) and male **b**). Large number of DE-TE-transcripts were only identified in one exposure for both female and male. The dot represented the different exposures. The dots connected by line indicated the number of DE-TE-transcripts shared across multiple exposures. **c**) Fraction of DE-TE-transcripts identified in both female and male separately for different exposures. only about 4% to 16% of DE-TE-transcripts were shared between female and male in across those exposures. **d**) Biology processes based on enriched TE subfamilies that derived protein-coding and retained-intron transcripts.

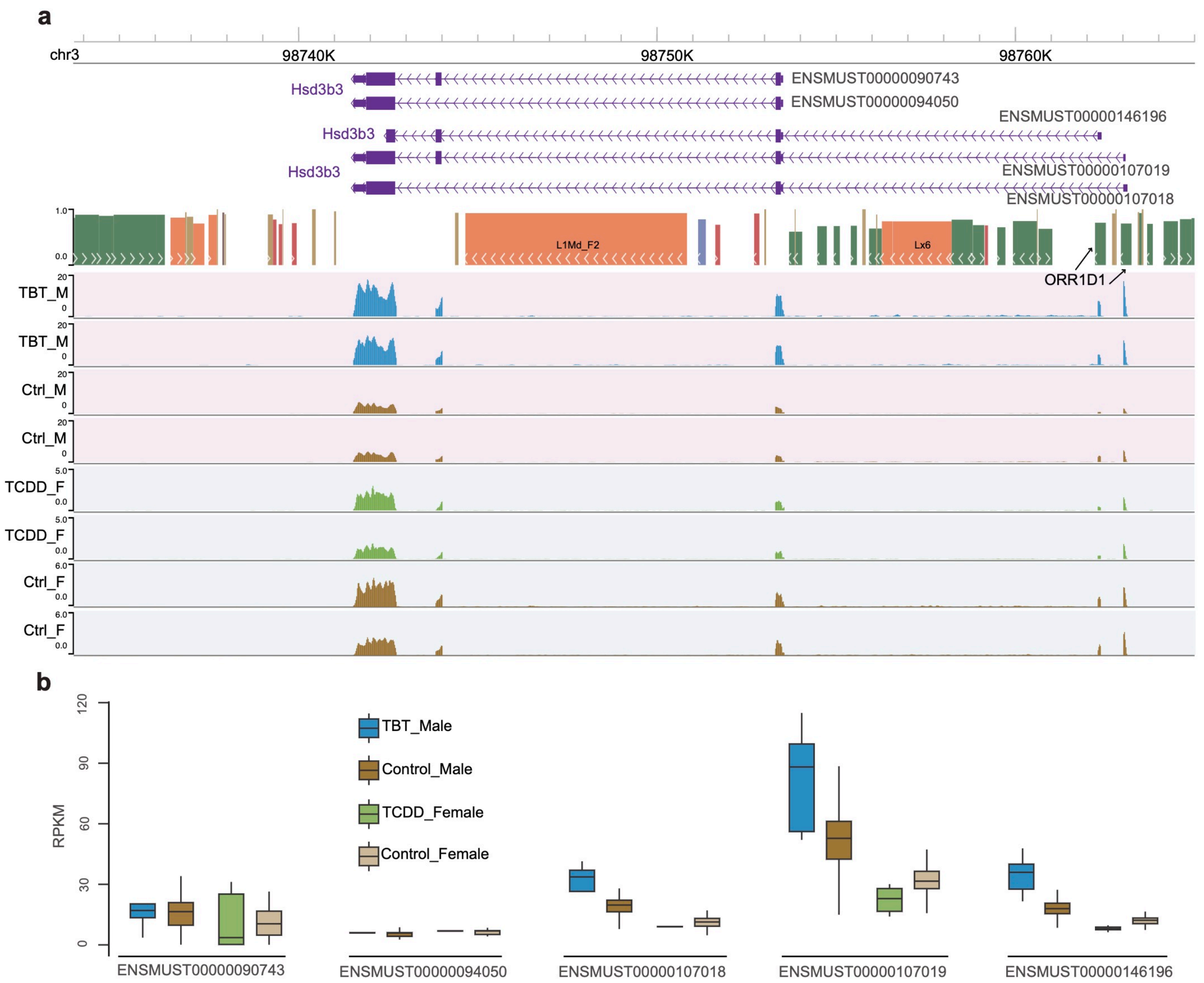


**S-figure 6.** Two ORR1D1 elements derived two alternative start sites of 3 isoforms of *Hsd3b3* gene. **a**) RNA-seq tracks of *Hsd3b3* gene from TBT, TCDD and control samples. b) Expression of transcripts of *Hsd3b3* gene. The TE derived isoforms were induced by TBT in male but depressed by TCDD and two BPA exposures in female. Meanwhile, the expression of those isoforms showed sex difference in control samples.

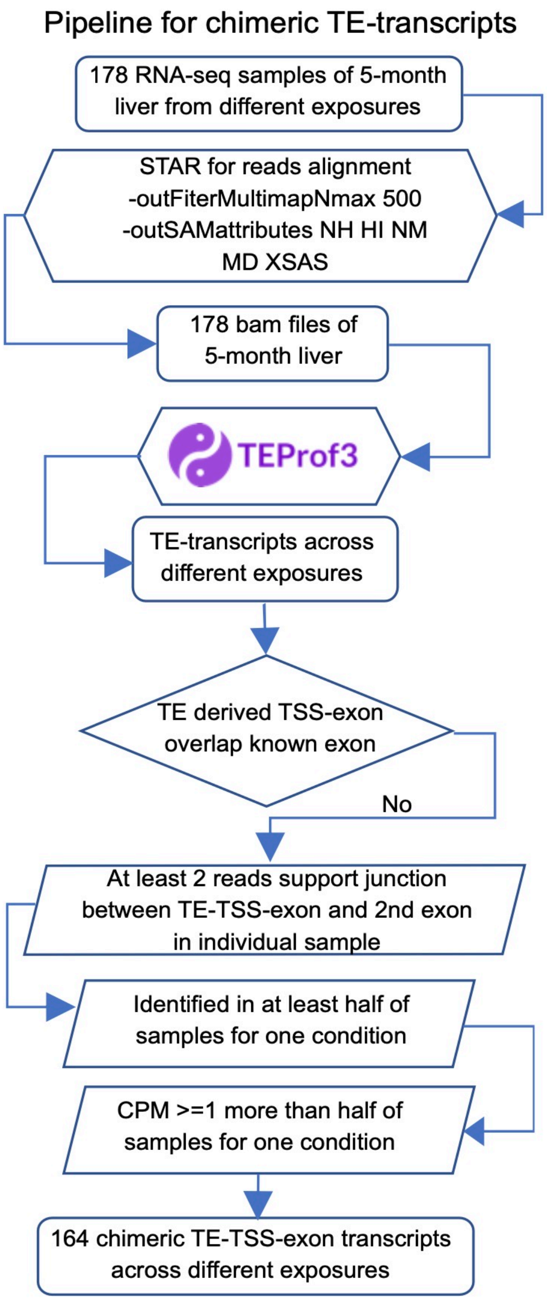


**S-figure 7.** Analysis pipeline to discover the TE derived chimeric transcripts (TE-chimeric-transcripts) induced by environmental exposures. STAR was used for reads alignment of RNA-seq data and TEProf3 was used to identify the TE derived transcripts. Total 164 TE-chimeric-transcripts were identified across different exposures that TE derived first exon did not overlap any known exons with at least 2 reads supporting junction between 1$^{st}$ and 2$^{nd}$ exon, and with CPM >=1 in more than half of samples for one condition.

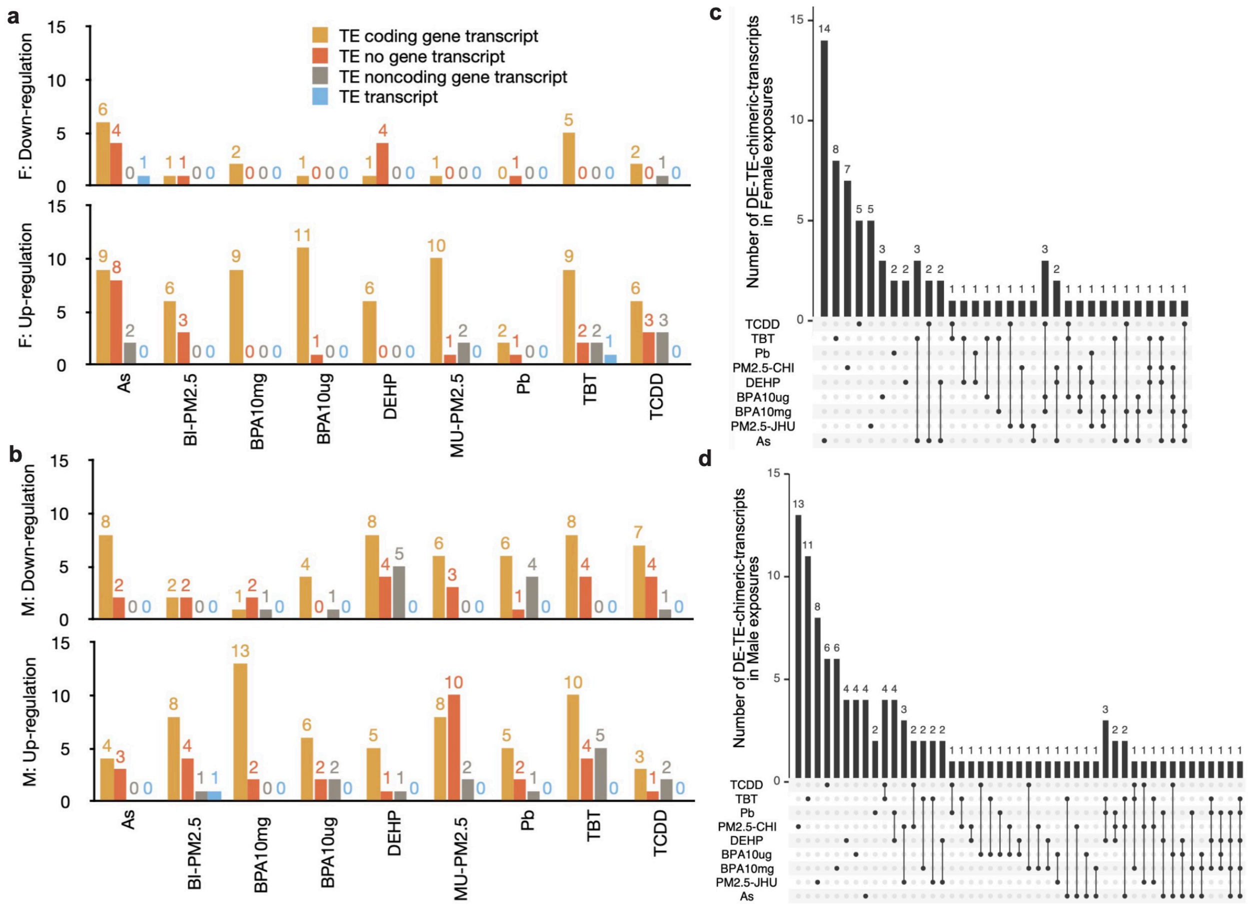


**S-figure 8.** Differentially expressed TE-chimeric-transcripts (DE-TE-chimeric-transcripts). Number of up and down-regulated DE-TE-chimeric transcripts of 4 different types across exposures in **a)** female and **b)** male. Large number of DE-TE-chimeric transcripts belonged to TE coding gene transcript and TE no gene transcript. Colors: different types. Distribution of DE-TE-chimeric transcripts across different exposures separately for **c)** female and **d)** male. Many DE-TE-chimeric transcripts were specific in one exposure for both female and male. The dots connected by line indicated number of DE-TE-chimeric transcripts shared by multiple exposures.

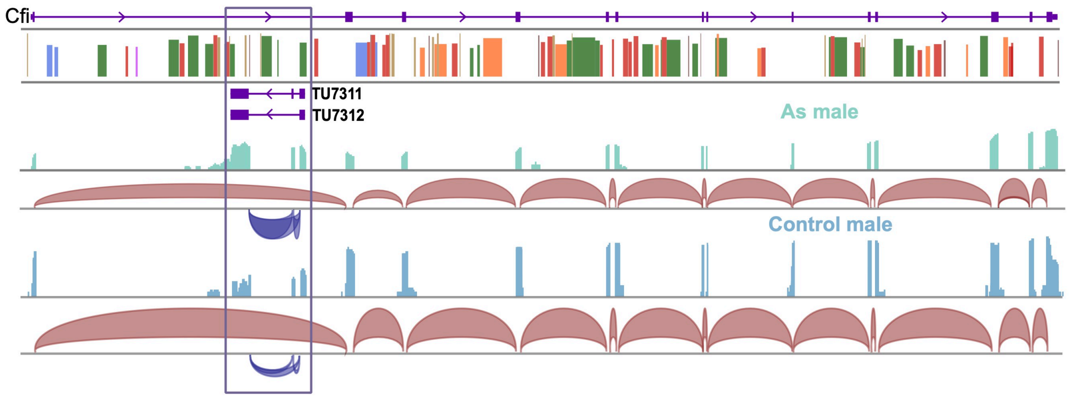


**S-figure 9.** RMER15 derived TE no gene transcripts (*TU7311* and *TU7312*) located in the intron of Cfi gene that was up regulated in As male comparing to control.
